# Principal component analysis-based unsupervised feature extraction applied to single-cell gene expression analysis^1^

**DOI:** 10.1101/312892

**Authors:** Y-h. Taguchi

**Affiliations:** Department of Physics, Chuo University, Tokyo 112-8551, Japan

**Keywords:** principal component analysis, feature selection, embryonic brain development

## Abstract

Due to missed sample labeling, unsupervised feature selection during single-cell (sc) RNA-seq can identify critical genes under the experimental conditions considered. In this paper, we applied principal component analysis (PCA)-based unsupervised feature extraction (FE) to identify biologically relevant genes from mouse and human embryonic brain development expression profiles retrieved by scRNA-seq. When evaluating the biological relevance of selected genes by various enrichment analyses, the PCA-based unsupervised FE outperformed conventional unsupervised approaches that select highly variable genes as well as bimodal genes in addition to the recently proposed dpFeature.

## 1 Introduction

Single-cell analysis is a newly developed high-throughput technology that enables us to identify gene expression profiles of individual genes. There is a critical difference between single-cell analysis and conventional tissue-specific analysis; tissue samples are labeled distinctively (e.g., patients vs healthy controls) while single-cell samples are not always. Inevitably, we need an unsupervised methodology, such as highly variable genes [1, 2] and bimodal genes [3] or recently proposed dpFeature [4]. Highly variable genes are able to select genes that can discriminate the underlying cluster structure depicted by unsupervised clustering, namely tSNE [5]. In contrast, bimodal genes are selected because unimodal genes are unlikely to be expressed distinctly between multiple classes, e.g., healthy controls and patients. While the combination of tSNE and highly variable genes or bimodal genes approach is often employed and empirically successful, biological validation of selected genes is rarely addressed. Generally, very few studies have evaluated multiple gene selection procedures for single-cell RNA-seq. The purpose of this paper is to compare multiple gene selection procedures and to identify the best method.

Principal component analysis (PCA)-based unsupervised feature extraction (FE) has been previously shown to be an effective method to investigate tissue-specific gene expression profiles [6–28]. In this paper, PCA-based unsupervised FE was applied to single-cell gene expression analysis. In addition, its effectiveness was evaluated from the biological point of view and compared to conventional approaches, including the highly variable genes approach as well as the bimodal genes approach.

## 2 Materials and Methods

### 2.1 Gene expression

Gene expression profiles used in this study were downloaded from the Gene Expression Omnibus (GEO) database under the GEO ID GSE76381. Specifically, the files named “GSE76381_EmbryoMoleculeCounts.cef.txt.gz” (for human) and “GSE76381_MouseEmbryoMoleculeCounts.cef.txt.gz” (for mouse) were downloaded. Gene expression profiles were standardized such that each sample had zero mean and unit standard deviation. That is, when *x*_*ij*_ represented the expression of *i*th gene in *j*th sample, Σ_*i*_ *x*_*ij*_ = 0 and 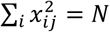 where *N* represented the number of genes. These two gene expression profiles were generated from single-cell RNA-seq datasets that represented the following: human embryo ventral midbrain cells between 6 and 11 weeks of gestation, mouse ventral midbrain cells at six developmental stages between E11.5 to E18.5, Th+ neurons at P19–P27, and FACS-sorted putative dopaminergic neurons at P28–P56 from Slc6a3-Cre/tdTomato mice.

### 2.2 PCA-based unsupervised FE

Suppose that matrix *X* has element *x*_ij_ representing the gene expression of *i*th gene of *j*th sample, then the *k*th PC score attributed to ith gene *u*_*ki*_ can be computed as *i*th element of *k*th eigen vector of Gram matrix *XX*^*T*^ as

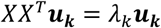

*k*th PC loading attributed to *j*th sample, *v*_*kj*_, can be obtained by ***v***_***k***_ = *X*^*T*^***u***_*k*_ since

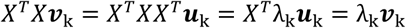

Initially, PC loading attributed to samples were identified, which were coincident with distinction between considered class labels attributed to the samples. Since single-cell RNA-seq (scRNA-seq) lacks sample labeling, the first *k* PCs were employed. Subsequently, assuming multiple Gaussian distribution to PC scores, *P*-values were attributed to gene *i* using χ^2^ distribution,

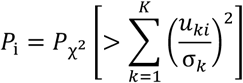

where σ_k_ represented standard deviation and *P*_χ^2^_ [> *x*] represented the cumulative probability of χ^2^ distribution that the argument was larger than *x*. The summation was taken over the selected first *k* PC scores. Obtained *P*-values were adjusted using Benjamini and Hochberg (BH) criterion [29] and genes associated with adjusted *P*-values less than 0.01 were selected.

### 2.3 Enrichment analysis

In order to perform enrichment analyses, selected genes were uploaded to Enrichr [30], which included various enrichment analysis.

### 2.4 Highly variable genes

The procedure was performed as previously described [1], and a brief description is provided below. Suppose that *x*_ij_ represented gene expression of *i*th gene of *j*th sample, then the mean expression of *i*th gene was defined as

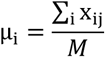

where M represented the number of samples. The standard deviation σ_i_ was defined as

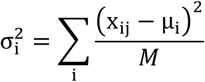

Following, the regression relation between σ_i_ and μ_i_ was assumed as

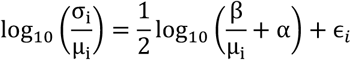

where α and β were the regression coefficients. Subsequently, P-value 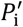 was attributed to ϵ_*i*_ assuming χ^2^ distribution as

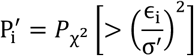

where σ′
was the standard deviation. Finally, genes *i* with an adjusted P-value less than 0.01 by BH criterion were selected as highly variable genes.

### 2.5 Bimodal genes

P-values attributed to genes that reject the null hypothesis (unimodal genes) were computed by the dip.test function in R using the default setting. Obtained P-values were adjusted by BH criterion, and genes associated with adjusted P-values less than 0.01 were selected.

### 2.6 dpFeature

dpFeature was performed using monocle package in R. More detailed instruction was in Supplementary Document.

## 3 Results

Applying PCA-based unsupervised FE to human and mouse embryonic brain developmental gene expression profiles, 116 genes for human (*k* = 2, i.e, the first two PC scores were used for gene selection) and 118 genes for mouse (*k* = 3, i.e., the first three PC scores were used for gene selection) were selected, respectively. Interestingly, 53 genes of the selected genes were common both in human and mouse samples. The large overlap between the two genes sets with highly restricted numbers of genes is not plausible to occur by chance; therefore, it is very likely that these selected genes play critical roles in embryonic midbrain development.

To validate the biological relevance of the selected genes, various enrichment analyses were applied using Enrichr. Table 1 shows an Enrichment analysis by Enrichr, “MGI Mammalian Phenotype 2017”, of the 118 genes selected in mice. Among the top five ranked terms, four were brain-related terms. As all terms described abnormal morphology, this is an expected result since fetal gene expression is often distinct from adults that lack fetal-specific gene expression. Table 2 and Table 3 show another Enrichment analysis by Enrichr, “Allen Brain Atlas down”, of the 116 genes selected in humans and 118 genes selected in mice, respectively. All genes were downregulated in brain regions. This is again reasonable since fetal genes expressed in the embryo is unlikely to be expressed in adult tissue. The lack of fetal brain-specific gene expression in the selected genes can also be seen in Table 4 and Table 5.

**Table 1.** Enrichment analysis by Enrichr, “MGI Mammalian Phenotype 2017”, of 118 selected genes in mice (Top 5 ranked terms)

| Term | Overlap | P-value | Adjusted P-value |
| --- | --- | --- | --- |
| MP:0000788_abnormal_cerebral_cortex_morphology | 7/145 | $2.45 \times 10^{-5}$ | $4.55 \times 10^{-5}$ |
| MP:0003651_abnormal_axon_extension | 5/48 | $9.18 \times 10^{-6}$ | $4.55 \times 10^{-3}$ |
| MP:0000812_abnormal_dentate_gyrus_morphology | 5/58 | $2.34 \times 10^{-5}$ | $4.55 \times 10^{-3}$ |
| MP:0000807_abnormal_hippocampus_morphology | 5/86 | $1.56 \times 10^{-4}$ | $2.04 \times 10^{-2}$ |
| MP:0000819_abnormal_olfactory_bulb_morphology | 4/48 | $1.83 \times 10^{-4}$ | $2.04 \times 10^{-2}$ |

**Table 2.** Enrichment analysis by Enrichr, “Allen Brain Atlas down”, of 116 selected genes in humans (Top 5 ranked terms)

| Term | Overlap | P-value | Adjusted P-value |
| --- | --- | --- | --- |
| periventricular stratum of cerebellar vermis | 18/300 | $1.33 \times 10^{-13}$ | $3.72 \times 10^{-11}$ |
| Simple lobule | 18/300 | $1.33 \times 10^{-13}$ | $3.72 \times 10^{-11}$ |
| Simple lobule, molecular layer | 18/300 | $1.33 \times 10^{-13}$ | $3.72 \times 10^{-11}$ |
| Simple lobule, granular layer | 18/300 | $1.33 \times 10^{-13}$ | $3.72 \times 10^{-11}$ |
| white matter of cerebellar vermis | 18/300 | $1.33 \times 10^{-13}$ | $3.72 \times 10^{-11}$ |

**Table 3.** Enrichment analysis by Enrichr, “Allen Brain Atlas down”, of 118 selected genes in mice (Top 5 ranked terms)

| Term | Overlap | P-value | Adjusted P-value |
| --- | --- | --- | --- |
| Pyramus (VIII), granular layer | 18/300 | $1.81 \times 10^{-13}$ | $4.66 \times 10^{-11}$ |
| Pyramus (VIII) | 18/300 | $1.81 \times 10^{-13}$ | $4.66 \times 10^{-11}$ |
| Pyramus (VIII), molecular layer | 18/300 | $1.81 \times 10^{-13}$ | $4.66 \times 10^{-11}$ |
| Paraflocculus, molecular layer | 18/300 | $1.81 \times 10^{-13}$ | $4.66 \times 10^{-11}$ |
| Cerebellar cortex | 18/300 | $1.81 \times 10^{-13}$ | $4.66 \times 10^{-11}$ |

**Table 4.**
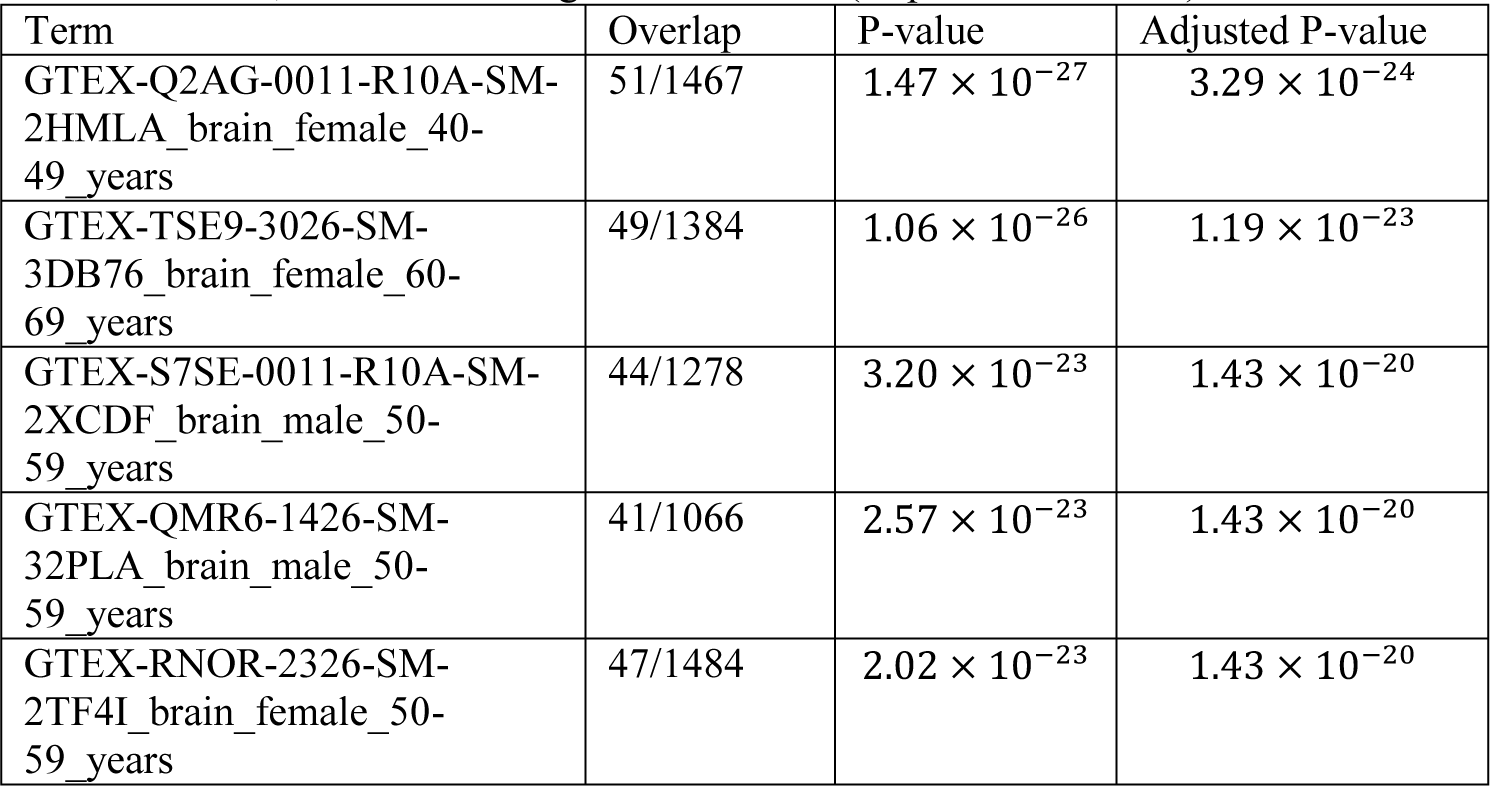
Enrichment analysis by Enrichr, “GTEx Tissue Sample Gene Expression Profiles down”, of 116 selected genes in humans (Top 5 ranked terms)

| Term | Overlap | P-value | Adjusted P-value |
| --- | --- | --- | --- |
| GTEx-Q2AG-0011-R10A-SM-2HMLA_brain_female_40-49_years | 51/1467 | $1.47 \times 10^{-27}$ | $3.29 \times 10^{-24}$ |
| GTEx-TSE9-3026-SM-3DB76_brain_female_60-69_years | 49/1384 | $1.06 \times 10^{-26}$ | $1.19 \times 10^{-23}$ |
| GTEx-S7SE-0011-R10A-SM-2XCDF_brain_male_50-59_years | 44/1278 | $3.20 \times 10^{-23}$ | $1.43 \times 10^{-20}$ |
| GTEx-QMR6-1426-SM-32PLA_brain_male_50-59_years | 41/1066 | $2.57 \times 10^{-23}$ | $1.43 \times 10^{-20}$ |
| GTEx-RNOR-2326-SM-2TF4I_brain_female_50-59_years | 47/1484 | $2.02 \times 10^{-23}$ | $1.43 \times 10^{-20}$ |

**Table 5.** Enrichment analysis by Enrichr, “GTEx Tissue Sample Gene Expression Profiles down”, of 118 selected genes in mice (Top 5 ranked terms)

| Term | Overlap | P-value | Adjusted P-value |
| --- | --- | --- | --- |
| GTEx-U8XE-0126-SM- | 15/376 | $6.13 \times 10^{-9}$ | $3.45 \times 10^{-6}$ |
| 4E3I3_testis_male_30-39_years |  |  |  |
| GTEX-X4XX-0011-R10B-SM-46MWO_brain_male_60-69_years | 23/938 | $5.25 \times 10^{-9}$ | $3.45 \times 10^{-6}$ |
| GTEX-U4B1-1526-SM-4DXSL_testis_male_40-49_years | 13/282 | $1.23 \times 10^{-8}$ | $3.71 \times 10^{-6}$ |
| GTEX-Q2AG-0011-R10A-SM-2HMLA_brain_female_40-49_years | 29/1467 | $5.11 \times 10^{-9}$ | $3.45 \times 10^{-6}$ |
| GTEX-RNOR-2326-SM-2TF4I_brain_female_50-59_years | 29/1484 | $6.62 \times 10^{-9}$ | $3.45 \times 10^{-6}$ |

While these above results are highly significant, they are also negative results. As such, we speculated whether these results could provide enough support and confidence for the selected genes. Therefore, in order to show positive results, gene expression in the embryonic brain was assessed. Subsequently, it was found that these genes were enriched in “Jensen TISSUES” by Enrichr. Table 6 shows that selected genes are enriched in the embryonic brains of humans and mice, respectively. Thus, confidence of the selected genes is supported by both negative and positive selection.

**Table 6.**
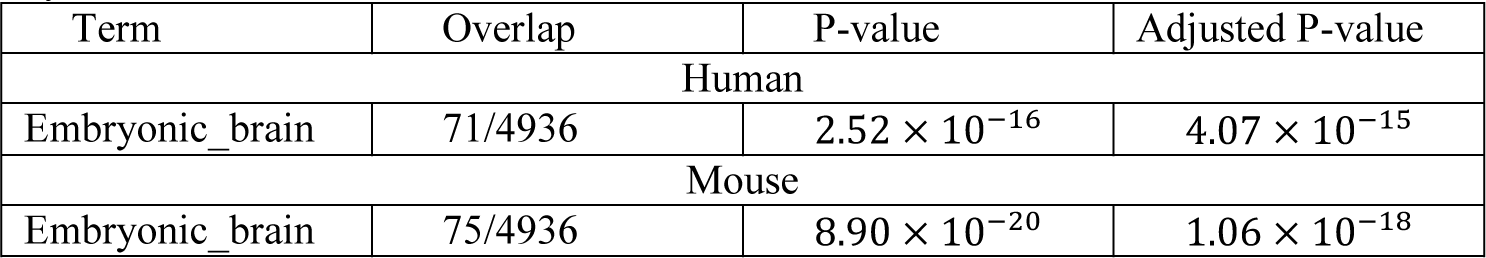
Selected gene enrichment in the embryonic brain of “Jensen TISSUES” by Enrichr

| Term | Overlap | P-value | Adjusted P-value |
| --- | --- | --- | --- |
| Human |  |  |  |
| Embryonic_brain | 71/4936 | $2.52 \times 10^{-16}$ | $4.07 \times 10^{-15}$ |
| Mouse |  |  |  |
| Embryonic_brain | 75/4936 | $8.90 \times 10^{-20}$ | $1.06 \times 10^{-18}$ |

We further sought the regulatory elements that can regulate the selected genes, as there are likely common regulatory elements if the selected genes are truly coexpressed. In order to perform this, we investigated “ENCODE and ChEA Consensus TFs from ChIP-X” by Enrichr for both human and mouse, respectively. Specifically, 42 TFs for humans and 23 TFs for mice were associated with adjusted *P*-values less than 0.01 (Table 7). Thus, they are likely co-regulated by these TFs. Moreover, most mouse TFs were also identified in humans (bold faces in Table 7). Therefore, it is very likely that we successfully identified common (species non-specific or conserved) TFs that regulate genes expression during embryonic brain development.

**Table 7.**
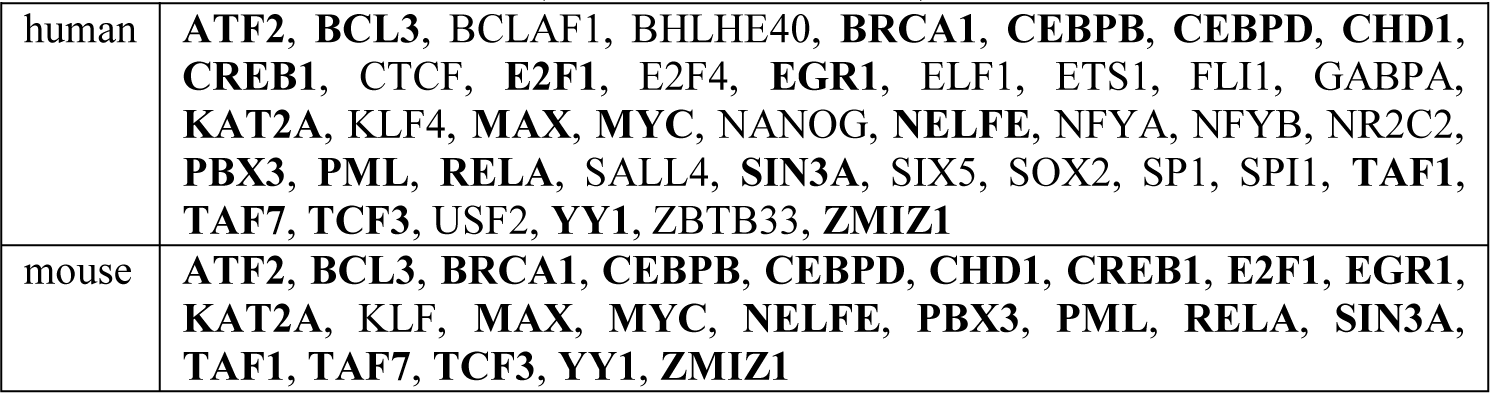
TF enrichment in “ENCODE and ChEA Consensus TFs from ChIP-X” by Enrichr for human and mouse (Bold TFs are common)

Additionally, we assessed whether the TF constructs functioned cooperatively. These TFs were uploaded to regnetwork server [31], and TF networks were identified as shown in Fig. 1. It is evident, even partially, that these TFs interact with each other.

**Fig. 1.**
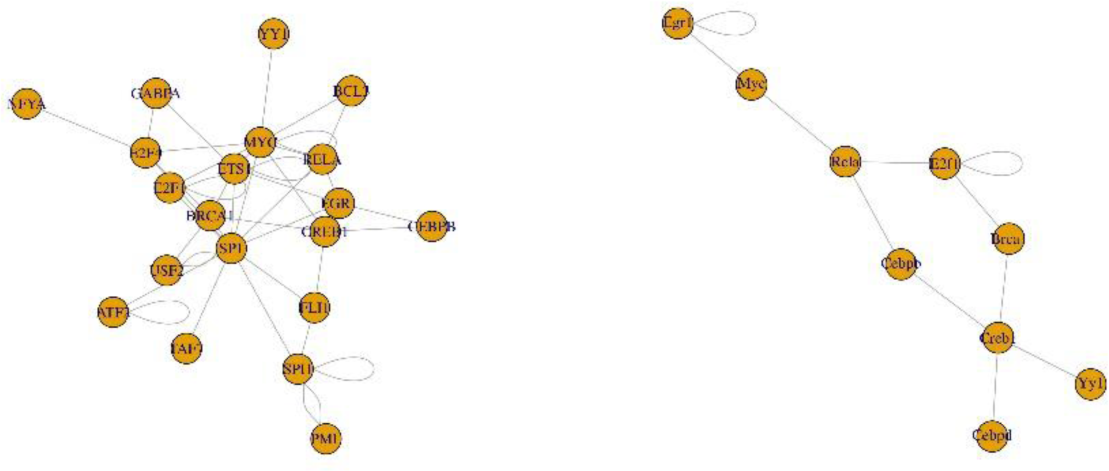
TF network identified by regnetworkweb for TFs in Table 7 (Left: human, right: mouse)

We also investigated whether the identified TFs were related to fetal brain development. TAF7 was reported to play a critical role in embryonic development [32]. KAT2A, ATF2 and TAF1 were also suggested to be included in brain development [33]. BRCA1 has been shown to play critical roles in brain development [34]. Additionally, CEBPD and CREB were reported to be related to brain disease [35, 36]. E2F1 was reported to be related to postnatal brain development [37] while functional EGR1 was found in the embryonic rat brain [38]. PML and SIN3A were also reported to be involved in brain development [39][40]. TCF3 has been shown to play a role in zebrafish brain development [41]. YY1 was also reported in brain development [42]. In conclusion, most of the selected genes in common (bold faces in Table 7) between humans and mice are related to brain development. Thus, the selection of genes is possibly reasonable.

In addition to the unconventional PCA-based unsupervised FE, we applied the widely used highly variable genes approach to the present data set and determined which strategy was more consistent with the enrichment analysis [1]. After applying the highly variable genes strategy, we obtained 168 genes for human and 171 genes for mouse. The numbers of selected genes were similar to those selected by PCA-based unsupervised FE. Additionally, there were 44 commonly selected genes between human and mouse. Thus, the highly variable genes method shows some efficacy.

However, after a detailed investigation, there were very few overlaps between genes selected by PCA-based unsupervised FE and highly variable genes (only four genes were commonly selected by PCA-based unsupervised FE and highly variable genes for both human and mouse). This result did not provide much confidence since genes selected by PCA-based unsupervised FE were demonstrated to be biologically reliable. However, there was still a possibility that highly variable genes were coincident to the enrichment analysis without significant overlap with genes selected by PCA-based unsupervised FE.

For confirmation, genes were uploaded to Enrichr. Accordingly, the results were substantially distinct from that given by PCA-based unsupervised FE. Specifically, the top five ranked enriched terms in “MGI Mammalian Phenotype 2017” for mouse did not include anything related to the brain, which is inferior to the results in Table 1. In contrast, no biological terms were significantly enriched in “Allen Brain Atlas down” for human, yet substantially large numbers of terms were enriched in mouse. This is inconsistent since the highly variable genes were substantially common between mouse and human; therefore, the inconsistency between human and mouse suggests that the selection of highly variable genes might be abiological. Subsequently, the “GTEx Tissue Sample Gene Expression Profiles down” was considered, and the top five ranked terms did not include anything related to brain, instead relations to skin and blood were found. This suggests that highly variable genes are not likely more biologically reliable than genes selected by PCA-based unsupervised FE. Finally, “Jensen TISSUES” was considered as was completed above (Table 6). Similarly, no brain-related terms were found to be significantly enriched in highly variable genes. TFs were also investigated to determine whether they could regulate highly variable genes. Nevertheless, “ENCODE and ChEA Consensus TFs from ChIP-X” only identified one TF whose target genes were significantly enriched in highly variable genes in human or mouse. As such, one TF identified is significantly less than that in Table 7. All of these suggest that highly variable genes are unlikely to be biologically more reliable than genes selected by PCA-based unsupervised FE.

In addition, we investigated bimodal genes as an alternative strategy (see Supplementary document). Genes associated with adjusted P-values less than 0.01 were 11344 and 10849 for human and mouse, respectively. Due to the large volume, it suggests that the bimodal genes approach has no ability to select a reasonable (restricted) number of genes. Nevertheless, in order to evaluate bimodal genes further, 200 top ranked (i.e., associated with smaller P-values) were intentionally selected. Specifically, of the 200 selected genes, only 21 genes were commonly selected between human and mouse, as compared to 53 and 44 commonly selected genes between human and mouse by PCA-based unsupervised FE and highly variable genes, respectively. Furthermore, there were no commonly selected genes between bimodal genes and PCA-based unsupervised FE. Enrichment analyses by Enricher for bimodal genes were also inferior to that of PCA-based unsupervised FE. “MGI Mammalian Phenotype 2017” of 200 bimodal genes selected for mouse included no terms associated with adjusted P-values less than 0.01 (Table S1). Top five ranked terms by “Allen Brain Atlas down” (Tables S2 and S3) were less significant than PCA-based unsupervised FE (Tables 2 and 3) as P-values were generally larger for bimodal genes. In addition, top five ranked terms by “GTEx Tissue Sample Gene Expression Profiles down” included no brain-related terms (Tables S4 and S5). Taken together, all of these suggests that bimodal genes are unlikely to be biologically more reliable than genes selected by PCA-based unsupervised FE.

Nevertheless, bimodal genes are slightly better than highly variable genes. Specifically, “Jensen TISSUES” by Enrichr included Embryonic_brain as a significant term (Table S6). In addition, “GTEx Tissue Sample Gene Expression Profiles down” identified 40 TFs associated with adjusted P-values less than 0.01 for human and mouse, among which as many as 30 TFs were commonly selected (Table S7). These TFs were also highly connected in the regnetworkweb (Figure S1).

Interestingly, among the 30 commonly selected TFs between human and mouse in Table S7, 11 TFs were also commonly selected between human and mouse in Table 7. When considering that no genes were commonly selected between top ranked 200 bimodal genes and genes selected by PCA-based unsupervised FE, the high number of commonly selected TFs between PCA-based unsupervised FE and bimodal genes suggests the robustness of TFs selected. Thus, as an overall evaluation PCA-based unsupervised FE is better than the other two.

Although we have also compared with the newly proposed approach, dpFeature, because of lack of space, it was included in Supplementary document. dpFeature could not select biologically more reliable genes than PCA based unsupervised FE, either.

While PCA is not a new technology and highly variable genes and bimodal genes are also not new concepts, the application to scRNA-seq is innovative and valuable. Therefore, even if methods themselves applied are not new, their application to new technology, scRNA-seq, can be innovative, especially if they have never been applied to scRNA-seq or were successful. Especially if PCA is more successful than even newly proposed approach, e.g., dpFeature.

## 4 Conclusions

We applied PCA-based unsupervised FE to gene expression profiles retrieved by scRNA-seq analysis. Since scRNA-seq primarily lacks the labeling of sample (each cell), an unsupervised approach is necessary. Genes selected by PCA-based unsupervised FE for human and mouse embryonic brain development were not only associated with numerous significant biological terms enrichment but also highly coincident between mice and humans. In contrast, the frequently employed highly variable genes approach as well as the bimodal genes approach or recently proposed dpFeature could not identify as many genes associated with significant biological terms enrichment as the PCA-based unsupervised FE achieved. Thus, PCA-based unsupervised FE is more favorable than the highly variable genes approach, the bimodal genes approach or dpFeature from the biological point of view.

## Supplementary materials

Supplementary materials are available at https://github.com/tagtag/SC.

Full list of genes, enrichment analyses for genes selected by PCA-based unsupervised FE, bimodal genes and dpFeature, supplementary document that includes supplementary tables and figure for bimodal gene analyses and dpFeature, R codes that identify genes selected by PCA-based unsupervised FE, highly variable genes, bimodal genes and dpFeature.

## Footnotes

1 Taguchi Y-h. (2018) Principal Component Analysis-Based Unsupervised Feature Extraction Applied to Single-Cell Gene Expression Analysis. In: Huang DS., Jo KH., Zhang XL. (eds) Intelligent Computing Theories and Application. ICIC 2018. Lecture Notes in Computer Science, vol 10955. Springer, Cham https://doi.org/10.1007/978-3-319-95933-7_90

## References

1. Chen, H.-I.H., Jin, Y., Huang, Y., Chen, Y.: Detection of high variability in gene expression from single-cell RNA-seq profiling. BMC Genomics. 17, 508 (2016).

2. Costa-Silva, J., Domingues, D., Lopes, F.M.: RNA-Seq differential expression analysis: An extended review and a software tool, (2017).

3. DeTomaso, D., Yosef, N.: FastProject: A tool for low-dimensional analysis of single-cell RNA-Seq data. BMC Bioinformatics. 17, (2016).

4. Qiu, X., Mao, Q., Tang, Y., Wang, L., Chawla, R., Pliner, H.A., Trapnell, C.: Reversed graph embedding resolves complex single-cell trajectories. Nat. Methods. 14, 979–982 (2017).

5. Maaten, L. Van Der, Hinton, G.: Visualizing Data using t-SNE. J. Mach. Learn. Res. 1. 620, 267–84 (2008).

6. Ishida, S., Umeyama, H., Iwadate, M., Taguchi, Y.H.: Bioinformatic Screening of Autoimmune Disease Genes and Protein Structure Prediction with FAMS for Drug Discovery. Protein Pept. Lett. 21, 828–839 (2014).

7. Taguchi, Y.-H.: microRNA-mRNA interaction identification in Wilms tumor using principalcomponent analysis based unsupervised feature extraction. In: 2016 IEEE 16th International Conference on Bioinformatics and Bioengineering (BIBE). pp. 71–78 (2016).

8. Murakami, Y., Kubo, S., Tamori, A., Itami, S., Kawamura, E., Iwaisako, K., Ikeda, K., Kawada, N., Ochiya, T., Taguchi, Y.-H.: Comprehensive analysis of transcriptome and metabolome analysis in Intrahepatic Cholangiocarcinoma and Hepatocellular Carcinoma. Sci. Rep. 5, 16294 (2015).

9. Taguchi, Y.-H.: Identification of More Feasible MicroRNA-mRNA Interactions within Multiple Cancers Using Principal Component Analysis Based Unsupervised Feature Extraction. Int. J. Mol. Sci. 17, 696 (2016).

10. Murakami, Y., Toyoda, H., Tanahashi, T., Tanaka, J., Kumada, T., Yoshioka, Y., Kosaka, N., Ochiya, T., Taguchi, Y. h.: Comprehensive miRNA Expression Analysis in Peripheral Blood Can Diagnose Liver Disease. PLoS One. 7, e48366 (2012).

11. Taguchi, Y.-H.: Identification of candidate drugs using tensor-decomposition-based unsupervised feature extraction in integrated analysis of gene expression between diseases and DrugMatrix datasets. Sci. Rep. 7, 13733 (2017).

12. Murakami, Y., Kubo, S., Tamori, A., Itami, S., Kawamura, E., Iwaisako, K., Ikeda, K., Kawada, N., Ochiya, T., Taguchi, Y.H.: Comprehensive analysis of transcriptome and metabolome analysis in Intrahepatic Cholangiocarcinoma and Hepatocellular Carcinoma. Sci Rep. 5, 16294 (2015).

13. Tamori, A., Murakami, Y., Kubo, S., Itami, S., Uchida-Kobayashi, S., Morikawa, H., Enomoto, M., Takemura, S., Tanahashi, T., Taguchi, Y.-H., Kawada, N.: MicroRNA expression in hepatocellular carcinoma after the eradication of chronic hepatitis virus C infection using interferon therapy. Hepatol. Res. 46, (2016).

14. Taguchi, Y.-H., Iwadate, M., Umeyama, H., Murakami, Y.: Principal component analysis based unsupervised feature extraction applied to bioinformatics analysis. Comput. Methods with Appl. Bioinforma. Anal. 153–182 (2017).

15. Taguchi, Y.H.: Principal Components Analysis Based Unsupervised Feature Extraction Applied to Gene Expression Analysis of Blood from Dengue Haemorrhagic Fever Patients. Sci. Rep. 7, 44016 (2017).

16. Taguchi, Y.-H., Iwadate, M., Umeyama, H., Murakami, Y., Okamoto, A.: Heuristic principal component analysis-based unsupervised feature extraction and its application to bioinformatics. (2014).

17. Taguchi, Y.-H.: Principal component analysis based unsupervised feature extraction applied to publicly available gene expression profiles provides new insights into the mechanisms of action of histone deacetylase inhibitors. Neuroepigenetics. 8, 1–18 (2016).

18. Taguchi, Y.-H., Murakami, Y.: Universal disease biomarker: can a fixed set of blood microRNAs diagnose multiple diseases? BMC Res. Notes. 7, 581 (2014).

19. Taguchi, Y.-H.: Principal component analysis based unsupervised feature extraction applied to budding yeast temporally periodic gene expression. BioData Min. 9, 22 (2016).

20. Umeyama, H., Iwadate, M., Taguchi, Y.-H.: TINAGL1 and B3GALNT1 are potential therapy target genes to suppress metastasis in non-small cell lung cancer. BMC Genomics. 15, S2 (2014).

21. Taguchi, Y.H., Murakami, Y.: Principal component analysis based feature extraction approach to identify circulating microRNA biomarkers. PLoS One. 8, e66714 (2013).

22. Taguchi, Y.-H., Wang, H.: Genetic Association between Amyotrophic Lateral Sclerosis and Cancer. Genes (Basel). 8, 243 (2017).

23. Taguchi, Y.-H., Iwadate, M., Umeyama, H.: SFRP1 is a possible candidate for epigenetic therapy in non-small cell lung cancer. BMC Med. Genomics. 9, (2016).

24. Taguchi, Y.-h., Iwadate, M., Umeyama, H.: Principal component analysis-based unsupervised feature extraction applied to in silico drug discovery for posttraumatic stress disorder-mediated heart disease. BMC Bioinformatics. 16, 139 (2015).

25. Taguchi, Y.-H., Iwadate, M., Umeyama, H.: Heuristic principal component analysis-based unsupervised feature extraction and its application to gene expression analysis of amyotrophic lateral sclerosis data sets. In: Computational Intelligence in Bioinformatics and Computational Biology (CIBCB), 2015 IEEE Conference on. pp. 1–10 (2015).

26. Y-H. Taguchi, Hideaki Umeyama, Mitsuo Iwadate, Yoshiki Murakami, Akira Okamoto: Heuristic Principal Component Analysis-Based Unsupervised Feature Extraction and Its Application to Bioinformatics. In: Baoying Wang, Ruowang Li, and William Perrizo (eds.) Big Data Analytics in Bioinformatics and Healthcare. pp. 138–162. IGI global (2015).

27. Murakami, Y., Tanahashi, T., Okada, R., Toyoda, H., Kumada, T., Enomoto, M., Tamori, A., Kawada, N., Taguchi, Y.H., Azuma, T.: Comparison of hepatocellular carcinoma miRNA expression profiling as evaluated by next generation sequencing and microarray. PLoS One. 9, (2014).

28. Taguchi, Y.-H.: Integrative Analysis of Gene Expression and Promoter Methylation during Reprogramming of a Non-Small-Cell Lung Cancer Cell Line Using Principal Component Analysis-Based Unsupervised Feature Extraction. In: ICIC2014. pp. 445–455 (2014).

29. Benjamini, Y., Hochberg, Y.: Controlling the false discovery rate: a practical and powerful approach to multiple testing. J. R. Stat. Soc. B57, 289–300 (1995).

30. Kuleshov, M. V., Jones, M.R., Rouillard, A.D., Fernandez, N.F., Duan, Q., Wang, Z., Koplev, S., Jenkins, S.L., Jagodnik, K.M., Lachmann, A., McDermott, M.G., Monteiro, C.D., Gundersen, G.W., Ma’ayan, A.: Enrichr: a comprehensive gene set enrichment analysis web server 2016 update. Nucleic Acids Res. 44, W90–W97 (2016).

31. Liu, Z.-P., Wu, C., Miao, H., Wu, H.: RegNetwork: an integrated database of transcriptional and post-transcriptional regulatory networks in human and mouse. Database. 2015, bav095 (2015).

32. Gegonne, A., Tai, X., Zhang, J., Wu, G., Zhu, J., Yoshimoto, A., Hanson, J., Cultraro, C., Chen, Q., Guinter, T., Yang, Z., Hathcock, K., Singer, A., Rodriguez-canales, J., Tessarollo, L., Mackem, S., Meerzaman, D., Buetow, K., Singer, D.S.: The General Transcription Factor TAF7 Is Essential for Embryonic Development but Not Essential for the Survival or Differentiation of Mature T Cells. Mol. Cell. Biol. 32, 1984–1997 (2012).

33. Tapias, A., Wang, Z.Q.: Lysine Acetylation and Deacetylation in Brain Development and Neuropathies. Genomics, Proteomics Bioinforma. 15, 19–36 (2017).

34. Pao, G.M., Zhu, Q., Perez-Garcia, C.G., Chou, S.-J., Suh, H., Gage, F.H., O’Leary, D.D.M., Verma, I.M.: Role of BRCA1 in brain development. Proc. Natl. Acad. Sci. 111, E1240–E1248 (2014).

35. Sun, Y., Jia, L., Williams, M.T., Zamzow, M., Ran, H., Quinn, B., Aronow, B.J., Vorhees, C. V., Witte, D.P., Grabowski, G.A.: Temporal gene expression profiling reveals CEBPD as a candidate regulator of brain disease in prosaposin deficient mice. BMC Neurosci. 9, 1–20 (2008).

36. Mantamadiotis, T., Lemberger, T., Bleckmann, S.C., Kern, H., Kretz, O., Villalba, A.M., Tronche, F., Kellendonk, C., Gau, D., Kapfhammer, J., Otto, C., Schmid, W., Schütz, G.: Disruption of CREB function in brain leads to neurodegeneration. Nat. Genet. 31, 47–54 (2002).

37. Suzuki, D.E., Ariza, C.B., Porcionatto, M.A., Okamoto, O.K.: Upregulation of E2F1 in cerebellar neuroprogenitor cells and cell cycle arrest during postnatal brain development. Vitr. Cell. Dev. Biol. - Anim. 47, 492–499 (2011).

38. Wells, T., Rough, K., Carter, D.A.: Transcription Mapping of Embryonic Rat Brain Reveals EGR-1 Induction in SOX2+ Neural Progenitor Cells. Front. Mol. Neurosci. 4, 1–12 (2011).

39. Korb, E., Finkbeiner, S.: PML in the Brain: From Development to Degeneration. Front. Oncol. 3, 1–5 (2013).

40. Witteveen, J. S., Willemsen, M. H., Dombroski, T. C. D., van Bakel, N. H. M., Nillesen, W. M., van Hulten, J. A., Jansen, E. J. R., Verkaik, D., Veenstra-Knol, H. E., van Ravenswaaij-Arts, C. M. A., Wassink-Ruiter, J. S. K., Vincent, M., David, A., Le Cai, S.M.: Haploinsufficiency of MeCP2-interacting transcriptional co-repressor SIN3A causes mild intellectual disability by affecting the development of cortical integrity. Nat. Genet. 48, 877–87 (2016).

41. Dorsky, R.I.: Two tcf3 genes cooperate to pattern the zebrafish brain. Development. 130, 1937–1947 (2003).

42. Beagan, J.A., Duong, M.T., Titus, K.R., Zhou, L., Cao, Z., Ma, J., Lachanski, C. V., Gillis, D.R., Phillips-Cremins, J.E.: YY1 and CTCF orchestrate a 3D chromatin looping switch during early neural lineage commitment. Genome Res. 27, 1139–1152 (2017).

